# Characterisation of RNA guanine-7 methyltransferase (RNMT) using a small molecule approach

**DOI:** 10.1101/2024.10.03.614057

**Authors:** Lesley-Anne Pearson, Alain-Pierre Petit, Cesar Mendoza Martinez, Fiona Bellany, De Lin, Sarah Niven, Rachel Swift, Thomas Eadsforth, Paul Fyfe, Marilyn Paul, Vincent Postis, Xiao Hu, Victoria H. Cowling, David W Gray

## Abstract

The maturation of the RNA cap involving guanosine N-7 methylation, catalyzed by the HsRNMT (RNA guanine-7 methyltransferase)-RAM (RNA guanine-N7 methyltransferase activating subunit) complex, is currently under investigation as a novel strategy to combat PIK3CA mutant breast cancer. However, the development of effective drugs is hindered by a limited understanding of the enzyme’s mechanism and a lack of small molecule inhibitors. Following the elucidation of the *Hs*RNMT-RAM molecular mechanism, we report the biophysical characterization of two small molecule hits. Biophysics, biochemistry and structural biology confirm that both compounds bind competitively with cap and bind effectively to *Hs*RNMT-RAM in the presence of the co-product SAM, with a binding affinity (K_D_) of approximately 1 μM. This stabilisation of the enzyme-product complex results in uncompetitive inhibition. Finally, we describe the properties of the cap pocket and provided suggestions for further development of the tool compounds.

## Introduction

RNA polymerase II transcripts are modified at the 5’ end by the co-transcriptional addition of the RNA cap [1,2]. In mammals this structure is 7-methylguanosine linked by a 5’ to 5’ triphosphate to the first transcribed nucleotide. The first and second transcribed nucleotides can be further methylated by other methyltransferases [3,4]. The RNA cap is formed during the early stages of transcription as the nascent RNA emerges from the polymerase complex [1,4]. RNGTT (RNA guanylyltransferase and 5’ phosphatase) binds directly to the RNA pol II large subunit and catalyses addition of the guanosine cap to the nascent transcript, denoted G(5’)ppp(5’)X where X is the first transcribed nucleotide. RNMT (RNA guanine-7 methyltransferase) catalyses transfer of a methyl group from S-adenosylmethionine (SAM) to the cap intermediate to form the cap m7G(5’)ppp(X) and the by-product S-adenosylhomocysteine (SAH). RNMT (RNA guanine-7 methyltransferase) was identified based on homology to the yeast and viral homologues and *in vitro* activity [5,6]. This basic cap structure (cap 0, m7G(5’)ppp(X)) protects RNA from degradation by nucleases during synthesis and recruits factors involved in RNA processing and translation initiation [1,2]. RNMT has an activating co-factor RAM (RNMT-activating miniprotein) clamped at the interface of the catalytic domain and the modular lobe 416-456 [7]. This stabilization promotes SAM recruitment and induces a significant increase in the methyltransferase activity [8, 9] through formation of a series of interactions affecting the α -helix A and multiple active site residues [10]. RAM also has an RNA binding domain that recruits RNA to RNMT [9,11. RNMT-RAM is critical for gene expression [12,13]. In addition to its role catalysing RNA cap methylation during transcription and later, RNMT-RAM also has non-catalytic roles in transcription and potentially other processes [11,14]. Deletion of RNMT results in reduced proliferation and increased apoptosis, and deregulated expression of RNMT promotes cell transformation [15,16,17,18].

Several cellular signalling pathways regulate RNMT, by modulating expression of RNMT itself or activating subunit RAM, by altering its activity through post-translational modifications, or by controlling metabolism of SAH, the inhibitory by-product of methylation reactions [1, 19, 20, 21]. Regulation of RNMT accompanies or drives major cellular transitions and directs enhanced expression of specific sets of genes. For example, RNMT-RAM is upregulated during T cell activation when rapid cell growth and proliferation requires increased mRNA synthesis for adaptive immune responses [12]. Of particular importance is the enhanced production of TOP-RNAs which encode ribosomal proteins and other proteins involved in ribosome production. Conversely, during embryonic stem cell differentiation, RNMT-RAM is down-regulated which is required for repression of pluripotency-associated genes and morphological features of differentiation [19].

Consequently, RNMT is a target of interest in cancers and immune disorders. These diseases exhibit deregulated transcription, which requires a cap, and RNMT inhibition has been demonstrated to reduce proliferation and increase apoptosis in many cell lineages [8,16,21]. Certain oncogenic mutations increase dependency on the cap. Breast cancer cell lines with oncogenic PIK3CA mutations have enhanced sensitivity to RNMT inhibition [15]. Myc-dependent transcription and proliferation is dependent on RNA cap formation [22,23,24]. Recent analyses have linked RNMT expression to cancer outcomes [25,26].

Here we use biophysical, biochemical and structural biology approaches to build understanding of the mode of enzymatic action of RNMT. Further, by carrying out a diversity screening campaign, we describe two novel small molecule inhibitors of RNMT that may be developable into useful tools for interrogating the role of RNMT in different cell contexts.

## Results

In this manuscript, sinefungin is used as a S-Adenosyl-Methionine (SAM) mimetic. cap is GpppG except in crystallographic studies where a cap surrogate Guanosine 5’-[β,γ-imido]triphosphate (GMP-PnP) was applied. Methylated cap and S-Adenosyl-L-homocysteine (SAH) are the final products of the enzymatic process.

### Biophysical and biochemical characteristics of RNMT

To elucidate the cap binding mode, full length *Hs*RNMT in complex with activating subunit RAM (Avi-HsRNMT-RAM) was investigated by SPR. We first measured the kinetic and equilibrium binding constants (*K*_D_) for SAM (Figure 1A), SAH (Figure 1B), and sinefungin (Figure 1C).

**Figure 1.**
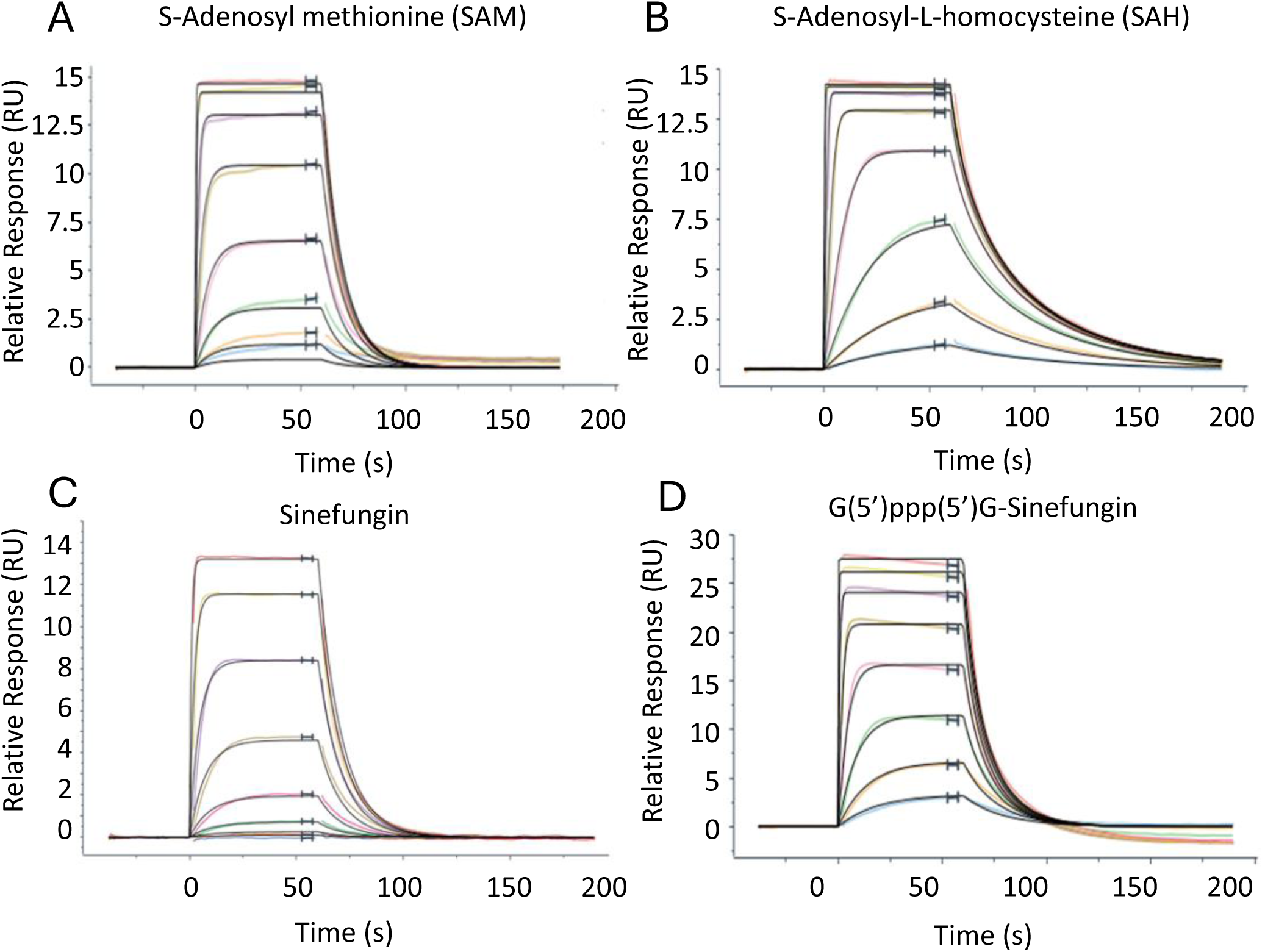
Sensograms derived from SPR binding experiments. All experiments were conducted with 8 concentrations in a 1:3 serial dilution. SAM - top concentration 5μM (A), SAH – top concentration 5μM (B), sinefungin – top concentration 1μM (C) or G(5’)ppp(5’)G-Sinefungin – top concentration 30μM were flowed over immobilized RNMT-RAM complexes. Data is representative of 2 experiments. Data was processed using Langmuir 1:1 model and is summarized in Table 1.

**Table 1.**
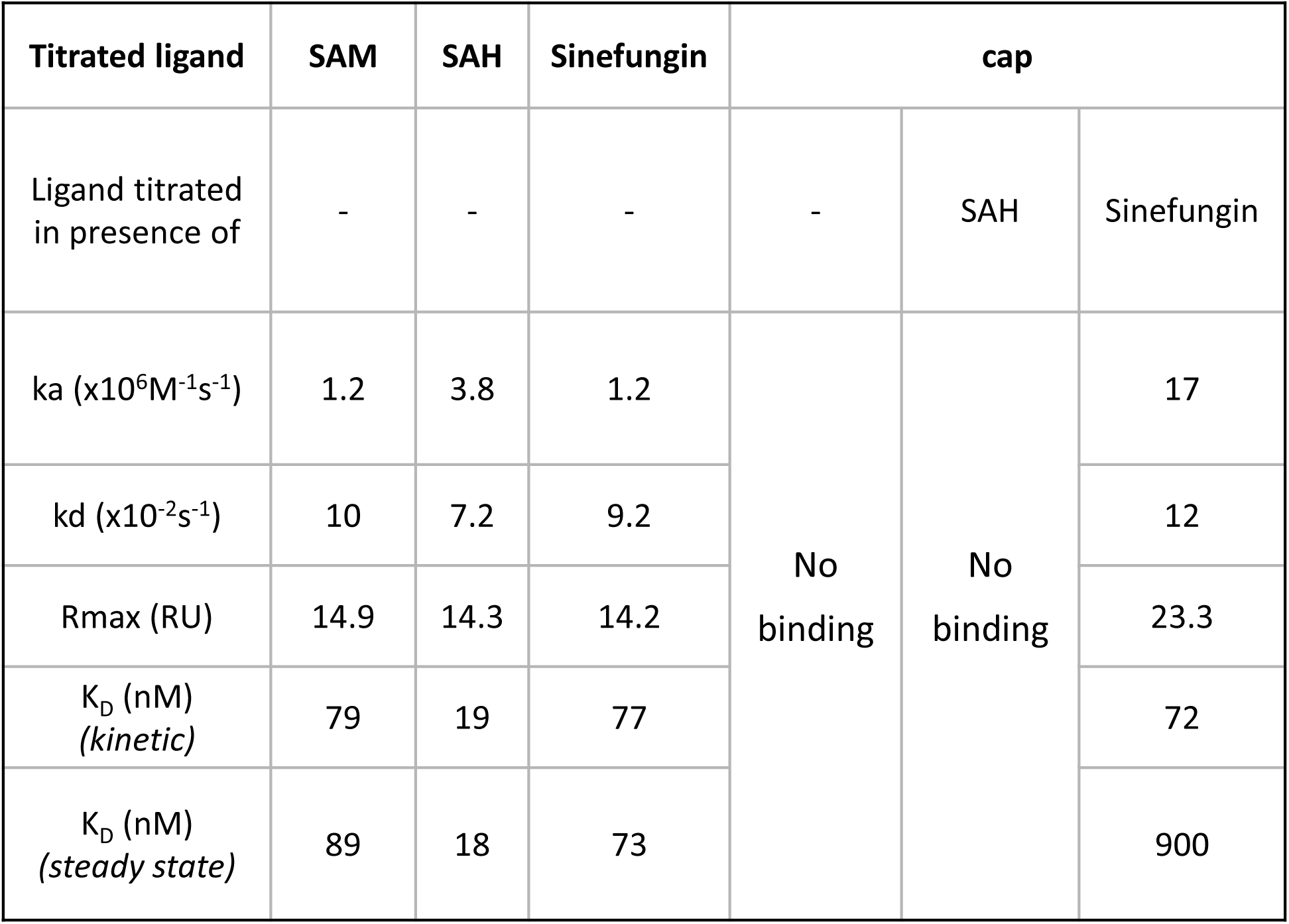
Summary of association (ka), dissociation (kd) and affinity (K_D_) estimates from the SPR assays. Rmax indictates the maximum binding capacity of the titrated fragment. Data were processed using a Langmuir 1:1 model.

No interaction of the cap with *Avi-HsRNMT-RAM* was detected in the absence of co-substrate or in the presence of SAH. The interaction of the cap with *Avi-HsRNMT-RAM* was only detected after addition of a saturating concentration (1µM) of sinefungin to the assay buffer (Figure 1D). These biophysical results are summarised in Table 1.

Altogether, these results suggest that *Hs*RNMT-RAM has an ordered binding mechanism with SAM binding first, followed by the cap. Furthermore, biophysical characterisations also suggest that the interaction of the cap with *Hs*RNMT-RAM was driven by the 5-amino group on sinefungin (or the SAM methyl group in a physiological condition) as SAH does not carry a comparable group at position 5 and no cap binding was detected. We hypothesise that in the absence of a 5-amino/methyl group, the cap guanine moiety is rotated by 180° in a clockwise direction, a consequence of the intra-molecular attractive force between the exocyclic -NH_2_ of the guanine moiety and the phosphates. This conformation would be detrimental to the cap binding since most of the interactions with *Hs*RNMT-RAM would be lost.

Michaelis-Menten parameters were determined for *Hs*RNMT-RAM using the RapidFire Mass Spec assay for cap (*K*_M_^cap^ = 3.2 µM [2.0-5.3 µM]), SAM (*K*_M_^SAM^ = 1.6 µM [1.3-2.1 µM]), and a mean *V*_Max_ 0.04 µM (0.04-0.05) SAH/min for both substrates (Figure 2).

**Figure 2.**
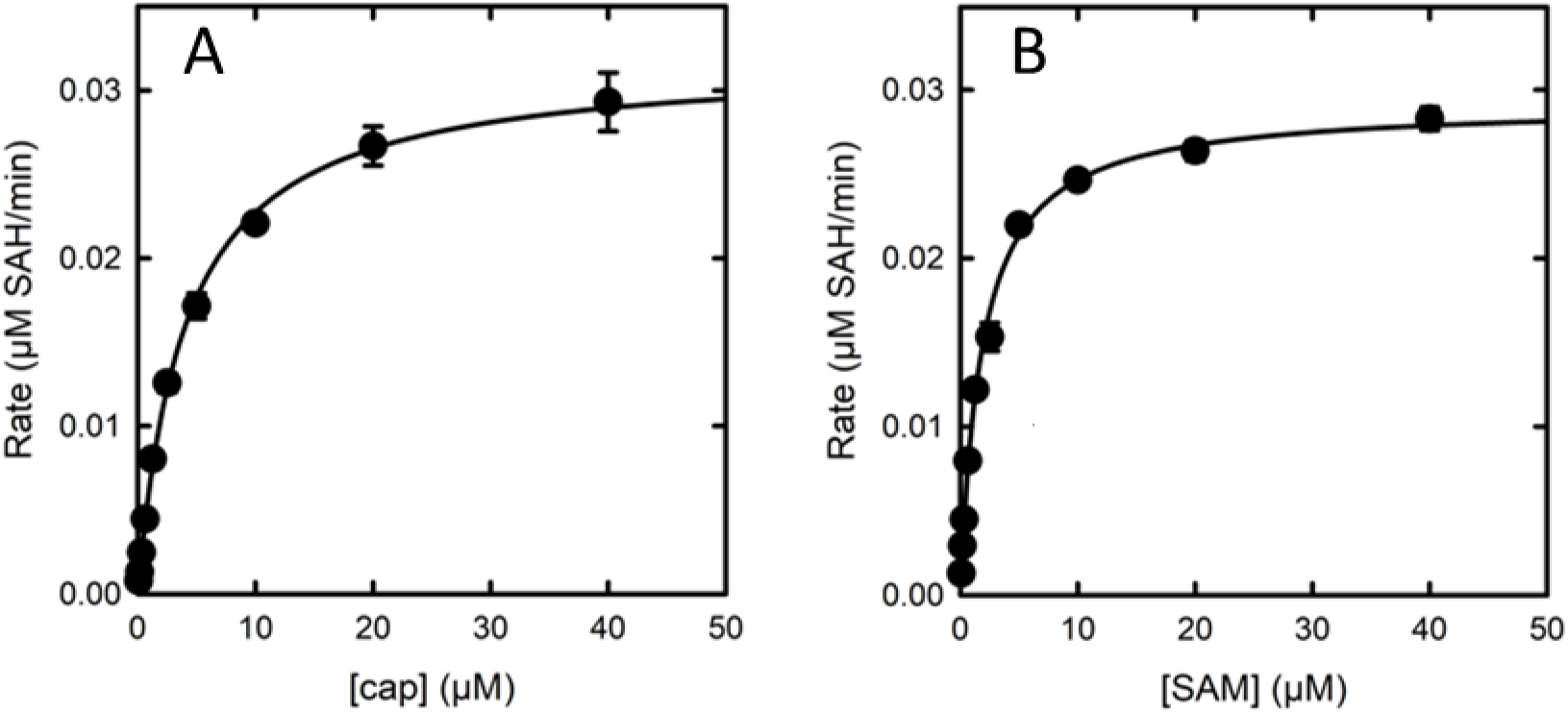
Michealis-Menten plots for HsRNMT-RAM substrates: cap (40 – 0.78 µM) (A) and SAM (40 – 0.078 µM) (B). Plots shown are representative examples from three experiments. Error bars are ± standard deviation from 4 technical replicates.

This data provides assay conditions suitable for characterisation of inhibitors. Sinefungin is a known competitive inhibitor of SAM in many enzymes [27] and this was confirmed for RNMT using the RapidFire assay (Table 1).

### Screening Campaign

Following the establishment of appropriate ‘balanced’ assay [28] a screening campaign was carried out on 48,806 diverse drug like compounds [29] from our internal compound library at a single concentration (10µM) using the MTAse Glo assay to measure enzymatic activity. Analysis of the single-point screen revealed a Normal distribution (Figure S1) with a mean inhibition of 4.6% and standard deviation of 15.4%. Using a cut-off of 50.8% (mean+3x standard deviation - 99.73% confidence assuming a normal hit distribution), the *Hs*RNMT-RAM assay resulted in a hit rate of 1.6%. The screen was robust with a mean Z’ [30] of 0.8 (±0.1) and mean signal:background of 7.7 (±3.6).

Two compounds with similar structures (Figure 3) were profiled in the orthogonal RapidFire assay.

**Figure 3.**
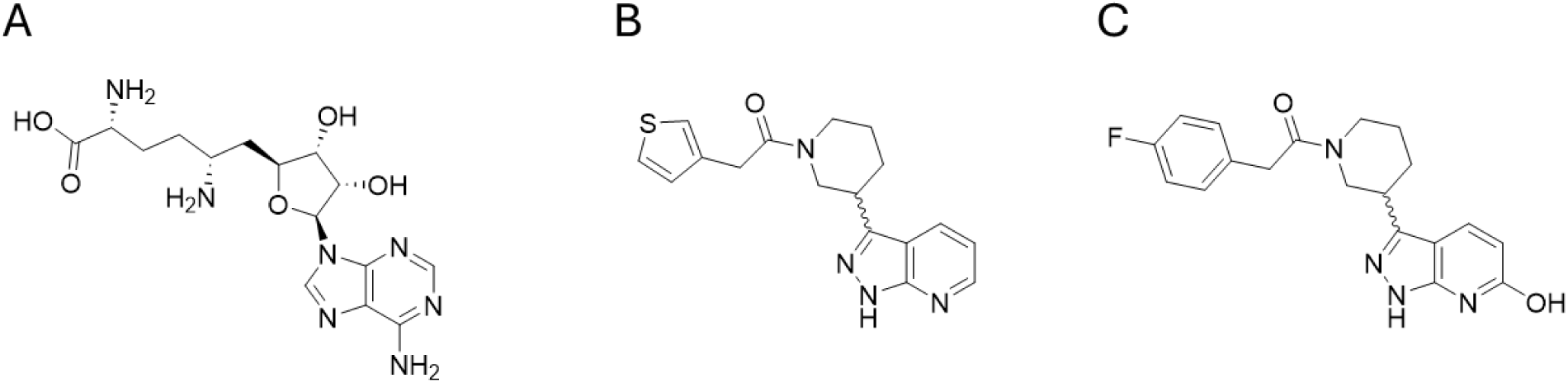
Chemical structures of (A) sinefungin. (B) DDD1060606 and (C) DDD1870799 DDD1060606 and DDD1870799 showed >80% inhibition of HsRNMT-RAM activity with determined pIC50s (negative logarithm of half-maximal inhibitory concentration [molar]) of 5.5±0.2 (IC50 3μM; Figure 4D and 5.0±0.2 (IC50 10μM; Figure 4G), respectively.

### Mode of Inhibition

Mode of inhibition experiments were performed for sinefungin, DDD1060606 and DDD1870799 using the RapidFire assay. While sinefungin showed competitive inhibition with SAM (Figure 4C) and non-competitive inhibition with cap (Figure 4B) as was anticipated, both small molecule inhibitors demonstrated uncompetitive inhibition profiles against both cap and SAM. (Figures 4E, F, H, I). The data from these experiments is summarised in Table 2.

**Figure 4.**
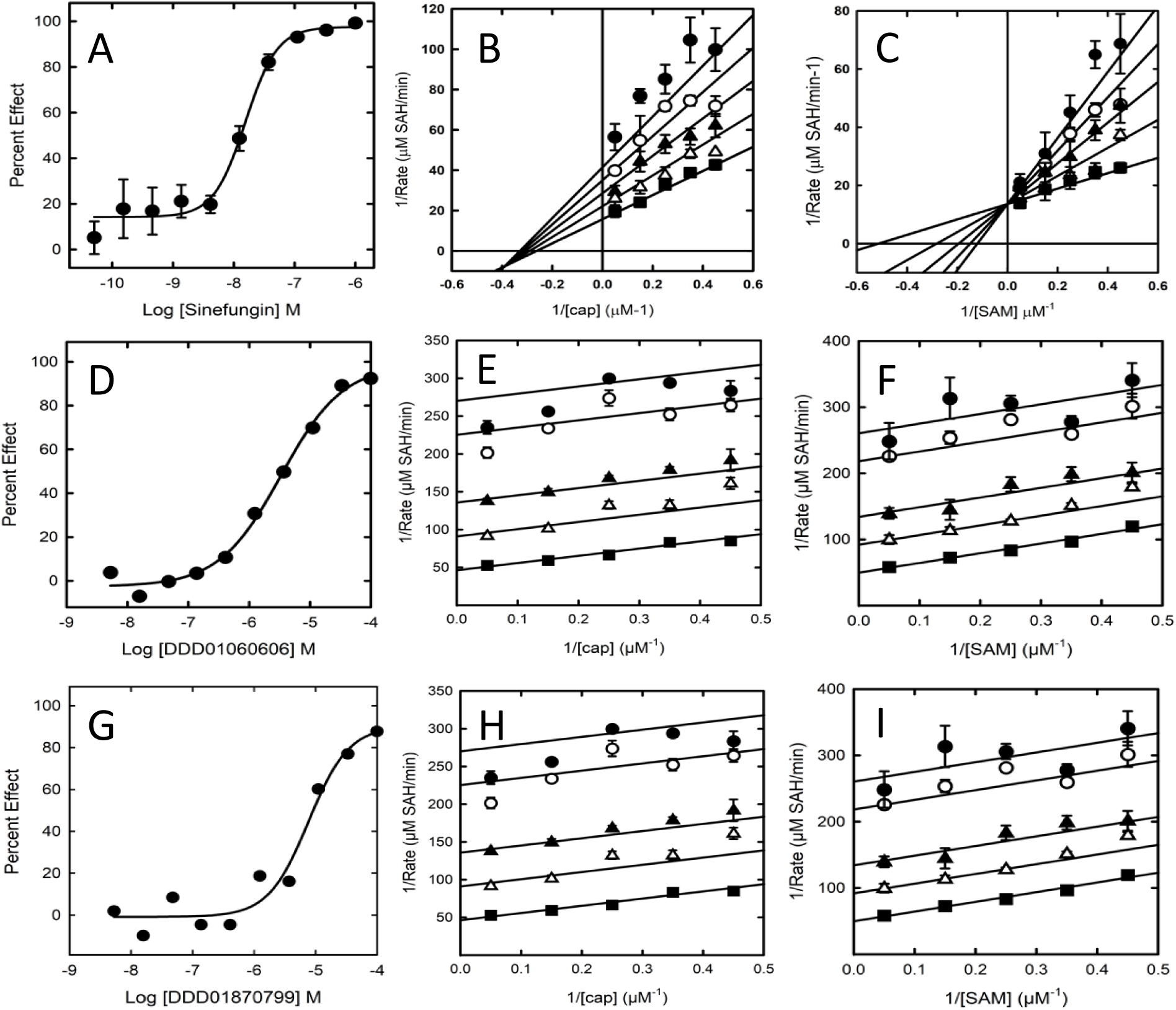
Characterisation of inhibition by sinefungin, DDD1060606 and DDD1870799. (A), (B) and (C) are the inhibition curve and Lineweaver-Burk plots against cap and SAM, respectively for sinefungin. (D), (E) and (F) are the inhibition curve and Lineweaver-Burk plots against cap and SAM, respectively for DDD1060606. (G), (H) and (I) are the inhibition curve and Lineweaver-Burk plots against cap and SAM, respectively for DDD1870799. Inhibition curves and Lineweaver-Burk plots are representative of 3 experiments. Error bars are standard deviation from 3 technical replicates

**Table 2.** Summary of mode of inhibition and key inhibition constants for sinefungin, DDD1060606 and DDD1870799. Data is presented as K_i_ (95% CI).

| Compound | K<sub>i</sub> cap | K<sub>i</sub> SAM |
| --- | --- | --- |
| sinefungin | Non-competitive<br>104.2 nM (92.2-117.8 nM) | Competitive<br>6.8 nM (3.1-14.9 nM) |
| DDD1060606 | Uncompetitive<br>1.6 μM (0.4-5.7 μM) | Uncompetitive<br>1.6 μM (0.4-5.7 μM) |
| DDD1870799 | Uncompetitive<br>3.3 μM (2.4-4.6 μM) | Uncompetitive<br>3.4 μM (2.8-4.2 μM) |

### Compound binding analysis

We used SPR to investigate the co-factor requirements to permit the binding of the inhibitors to *Hs*RNMT-RAM. No binding was observed in absence of co-ligand or in presence of SAM whereas high affinity binding (K_D_=1.6 µM and 2.5 µM (Figure 5) was measured for DDD1060606 and DDD1870799, respectively when the buffer was supplemented with SAH. This suggests that the ligands bind preferentially after the reaction has completed and SAM has been metabolised into SAH.

**Figure 5.**
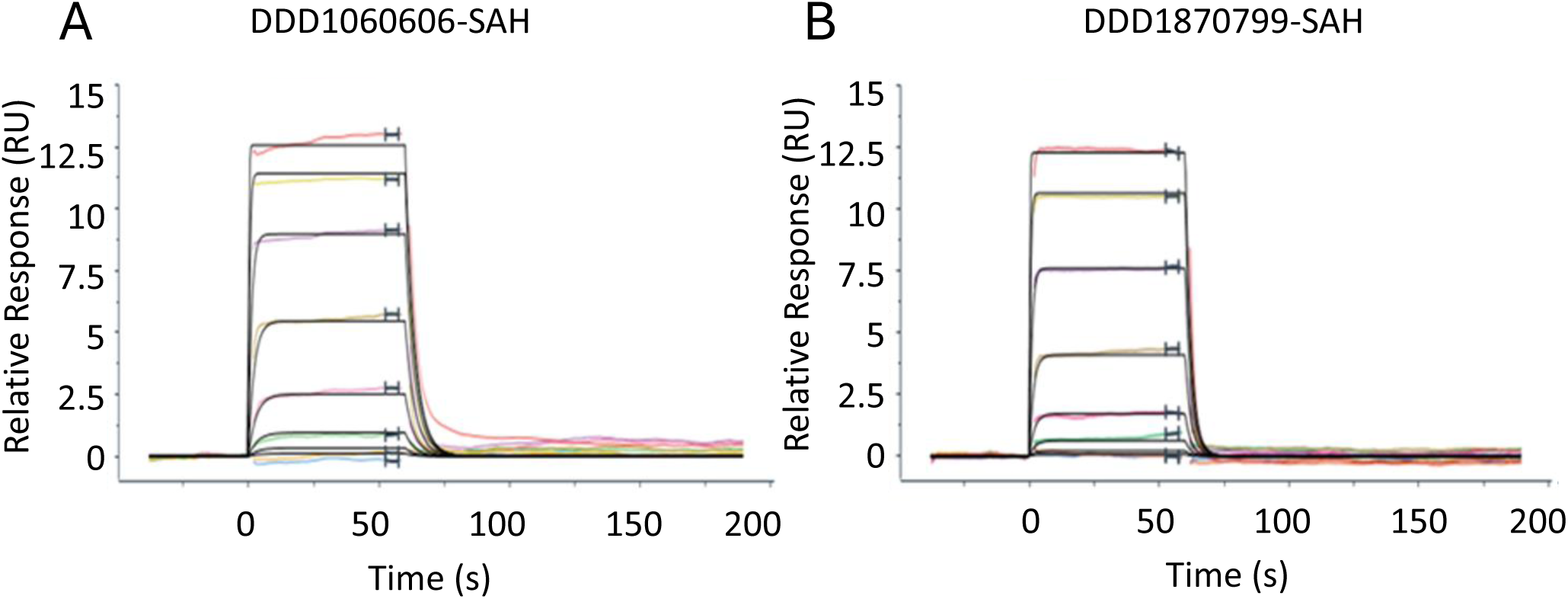
Sensograms derived from SPR binding experiments. All experiments were conducted with an 8 concentration in a 1:3 serial dilution. DDD1060606 – top concentration 30μM (A) or DDD1870799 – top concentration 30μM (B) were flowed over immobilized RNMT-RAM complexes in the presence of SAH (what concentration). Data is representative of 2 experiments. Data was processed using Langmuir 1:1 model and is summarized in Table 3.

**Table 3.**
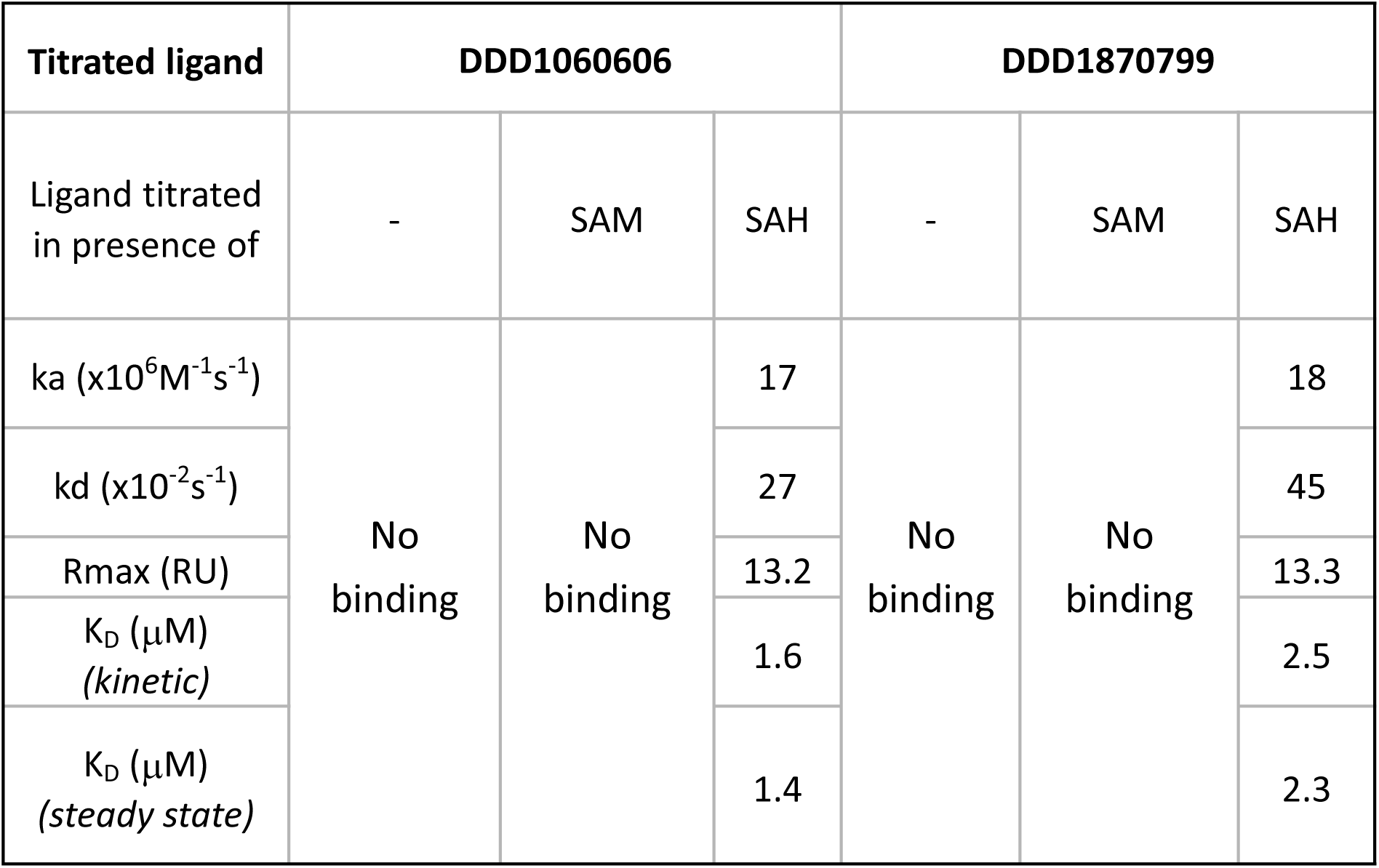
Summary of association (ka), dissociation (kd) and affinity (K_D_) estimates from the SPR assays. Rmax indictates the maximum binding capacity of the titrated fragment. Data were processed using a Langmuir 1:1 model.

The data from these experiments are summarised in Table 3.

### Crystallography

To improve the robustness of the crystallisation process, the lobe 416-455 of *Hs*RNMT catalytic domain was replaced by its homologous sequence region from the *Encephalitozoon cuniculi* (GLGC) guanine-7 cap methyltransferase (Q8SR66) identified from a pairwise sequence alignment. The 2 chains of the asymmetric unit for all structures are almost identical and can be well superposed (RMSD < 0.5 Å) to the structure of reference (5e8j) solved in complex with RAM [1] (Figure S2). This validates our hypothesis that the effect of this lobe substitution would be negligible on the RNMT-RAM structure.

### SAM/SAH binding site

*Hs*RNMT-Sinefungin-GMP-PnP, *Hs*RNMT-SAH-DDD1060606 and *Hs*RNMT-SAH-DDD1870799 co-complexes were solved respectively at 2.5, 2.0 and 2.4 Å in space group *P*2_1_ (Table S1).

In all 3 structures, SAH or sinefungin are observed in a hydrophobic pocket made by the α-helices 3 and 4 and the loops connecting the β-strand 1 to the α-helix B and the β-strands 3 and 4. SAH and sinefungin are stabilised in the active site through an identical network of hydrogen bonds, involving residues K180, G205, D227, D261, S262 and Q284 as well as 2 salt bridges with K180 (Figure S3). These results support the hypothesis that sinefungin binds in the SAM binding site in the presence or absence of the cap.

### cap binding analysis

The understanding of the cap binding mechanism is currently limited by the absence of a published *Hs*RNMT-cap structure and, lack of knowledge of the key residues involved in cap binding. This deficit hinders the development of potent and/or selective drugs targeting the cap pocket. Since attempts to crystallize RNMT with the G(5’)ppp(5’)G (cap) had been unsuccessful, we co-crystallized RNMT with the cap surrogate, Guanosine 5’-[β,γ-imido]triphosphate (GMP-PnP). GMP-PnP was confirmed as a cap competitive inhibitor with a Ki of 2.3 µM (0.9-5.5 µM) (Figure S4). Crystallisation of RNMT with GMP-PnP provides the first structural characterisation of a Class I methyltransferase in a complex with a cap analogue and sinefungin. The sinefungin molecule is well resolved in the electron density map (Figure S5) for both chains of the asymmetric unit and shares a common network of interaction to that observed for SAH binding to *Hs*RNMT (Figure 6A). GMP-PnP was successfully built in just one of the chains in the asymmetric unit and is located between the α-helix A and the β-strands 8 / 9. When co-crystallized with HsRNMT and sinefungin, the GMP-PnP guanine moiety is stabilized within the cap pocket by hydrogen bonds to residues H288, Y289, E370, Y467 and the sinefungin 5-amino group, as well as through a Pi-stacking interaction with F285. The GMP-PnP ribose makes two hydrogen bonds with N176 whereas the beta-gamma pyrophosphate group interacts with the polypeptide chain via two salt bridges to K208. Finally, the distance measured between the sinefungin 5-amino group and the guanine N7 (2.4 Å) alongside the angle formed by C5/5-NH2/N7 (140°) confirms GMP-PnP is observed in an ideal orientation to permit the methyl transfer, similar to the predicted model published [10].

**Figure 6.**
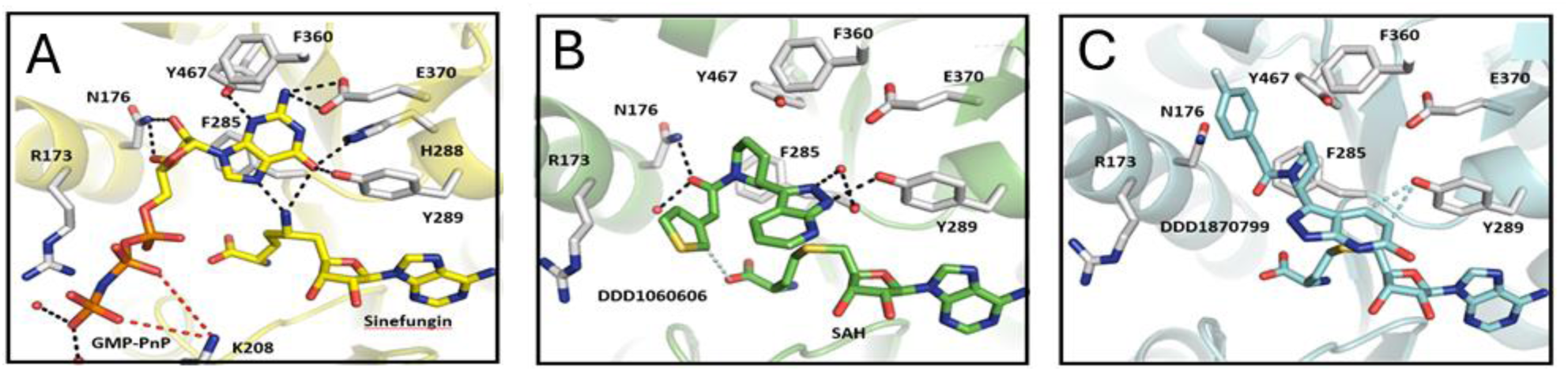
Structural analysis of the HsRNMT cap binding site. (A) Guanosine 5′-[β,γ-imido]triphosphate in complex with sinefungin, (B) DDD1060606 in complex with SAH and (C) DDD1870799 in complex with SAH. Residues within 4Å are drawn in sticks. Dash lines represent hydrogen bonds (black), salt bridges (red) and aromatic H-bonds (cyan).

The two inhibitors (DDD1060606 and DDD1870799) identified in the screening campaign were successfully co-crystallized with *Hs*RNMT in the presence of SAH. DDD1060606 (Figure 6B) is observed at the cap pocket and displays a mainly hydrophobic binding mode. Using the *Hs*RNMT-Sinefungin-GMP-PnP as a structure of reference, the pyrazolo-pyridine is rotated 180° and sits on top of the SAH homocysteine moiety, interacting with F285 and Y289 via two hydrogen bonds. The oxo-piperidine is located nearby the GMP-PnP ribose and is mainly stabilised by a hydrogen bond with N176. Finally, the thiophene near to the GMP-PnP alpha phosphoryl group is largely solvent exposed and is only stabilised by a single aromatic H-bond with SAH.

DDD1870799 (Figure 6C) is also observed in the cap pocket of RNMT. In the presented structure, the pyrazolo-pyridine interacts with Y289 via 2 aromatic H-bonds while the fluoro-benzene is buried in a hydrophobic pocket made by the residues including the main chain R173 as well as the side chains N176 and Y467.

### Active site characterisation

We determined DDD1060606/DDD1870799 enthalpic and entropic components of the free energy of binding to *Hs*RNMT-RAM to guide both compound selection and the optimisation process. We used cap as the reference ligand as it is widely accepted that ligands with more negative enthalpies of binding provide better starting points for lead optimization. The interaction of cap with *Hs*RNMT-RAM is enthalpically driven with an ΔH of −33.5 kcal/mol and net free energy change, ΔG of −8.4 kcal/mol. Furthermore, we determined the binding affinities for DDD1060606 and DDD1870799 to *Hs*RNMT-RAM. The resulting affinities (both approximately 3 μM) are comparable to the K_D_ measured by SPR in the previous experiments. A favourable entropic contribution Δ(TΔS) of −1.8 kcal/mol was observed for DDD1060606 and the total Gibbs free energies were similar for both ligands (ΔG = −7.5 kcal/mol). However, a significant gain in net enthalpic change ΔΔH of −1.9 kcal/mol was measured for DDD1870799 in comparison to DDD1060606 suggesting the interaction of DDD1870799 with *Hs*RNMT-RAM is more enthalpy driven and might result in more selective leads with fewer ADMET (Absorption, Distribution, Metabolism, Excretion and Toxicity) issues than DDD1060606 (Figure 7).

**Figure 7.**
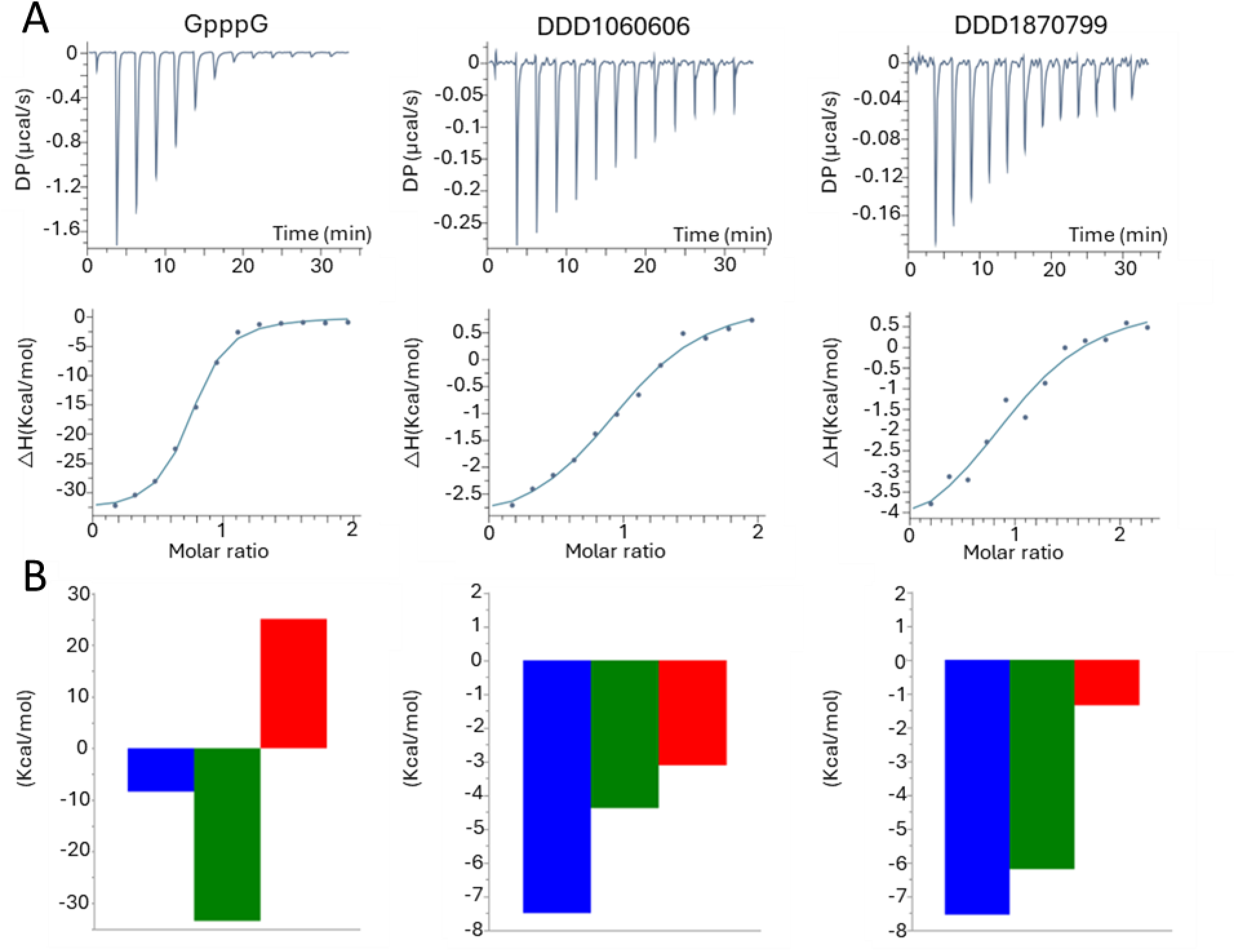
cap, DDD1060606 and DDD1870799 titrations into RNMT-RAM. (A) Raw data and integration of data, corrected for the heat of dilution. The line represents the least-squares fit to the single-site binding model by the ORIGIN program. (B) Thermodynamic signatures (ΔG° in blue, ΔH° in green, − TΔS° in red, all data in kJ/mol).

The data from this analysis is summarized in Table 4.

**Table 4.** Summary of mode of inhibition and key inhibition constants for sinefungin, DDD1060606 and DDD1870799. ITC titrations were performed in 25mM HEPES pH7.5, 150mM NaCl, 1mM TCEP supplemented with 10μM sinefungin or SAH at 298°K.

| Ligand | $\Delta G$<br>(kcal/mol) | $\Delta H$<br>(kcal/mol) | $-T\Delta S$<br>(kcal/mol) | N | $K_D$ ($\mu M$) |
| --- | --- | --- | --- | --- | --- |
| cap | -8.4 | -33.5 | 25.1 | 0.7 | 0.7 |
| DDD1060606 | -7.5 | -4.3 | -3.1 | 1.0 | 3.2 |
| DDD1870799 | -7.5 | -6.2 | -1.3 | 1.0 | 2.9 |

We then calculated the molecular interaction fields (MIFs) by FLAP [31] to identify the attributes that lead to the biological activity of the cap analogue and the 2 new chemical entities (NCE).

Figures 8A and B show the calculated FLAP maps for the cap analogue (GMP-PnP) binding to RNMT. The observed polar back pocket requires a hydrogen bond donor (guanine - position 2) to interact with E370 that is bordered by hydrophobic regions (pocket S1) while the position of the 3’O (ribose) coordinates well the requirement for a hydrogen bond acceptor to generate the hydrogen bond with N176. The DDD1060606 oxo-piperidine (Figure 8C), meanwhile, matches the hydrophobic and the hydrogen bond acceptor requirements introduced by F285 and N176 respectively (pocket S2). It is worth noting that neither the pyrazolo-pyridine or the thiophene can form favourable interactions with the protein, while the former sits above the thioether group of SAH. In comparison, the MIFs calculated for DDD1870799 (Figure 8D) suggest the fluoro-phenyl moiety is responsible for the ligand binding via the interaction with a small pocket (S3) of high hydrophobicity. This interaction, by displacement of disorganized water molecules [32] is likely to be responsible for the significant gain in net enthalpic change ΔΔH measured for DDD1870799. Finally, since the orientations observed for the NCE pyrazolo-pyridine groups do not appear to provide any advantage in comparison with guanine from the cap, the design of ligands with an improved affinity profile will have to take into consideration substituents that may be capable of stabilising the pyrazolo-pyridine in the S1 pocket.

**Figure 8.**
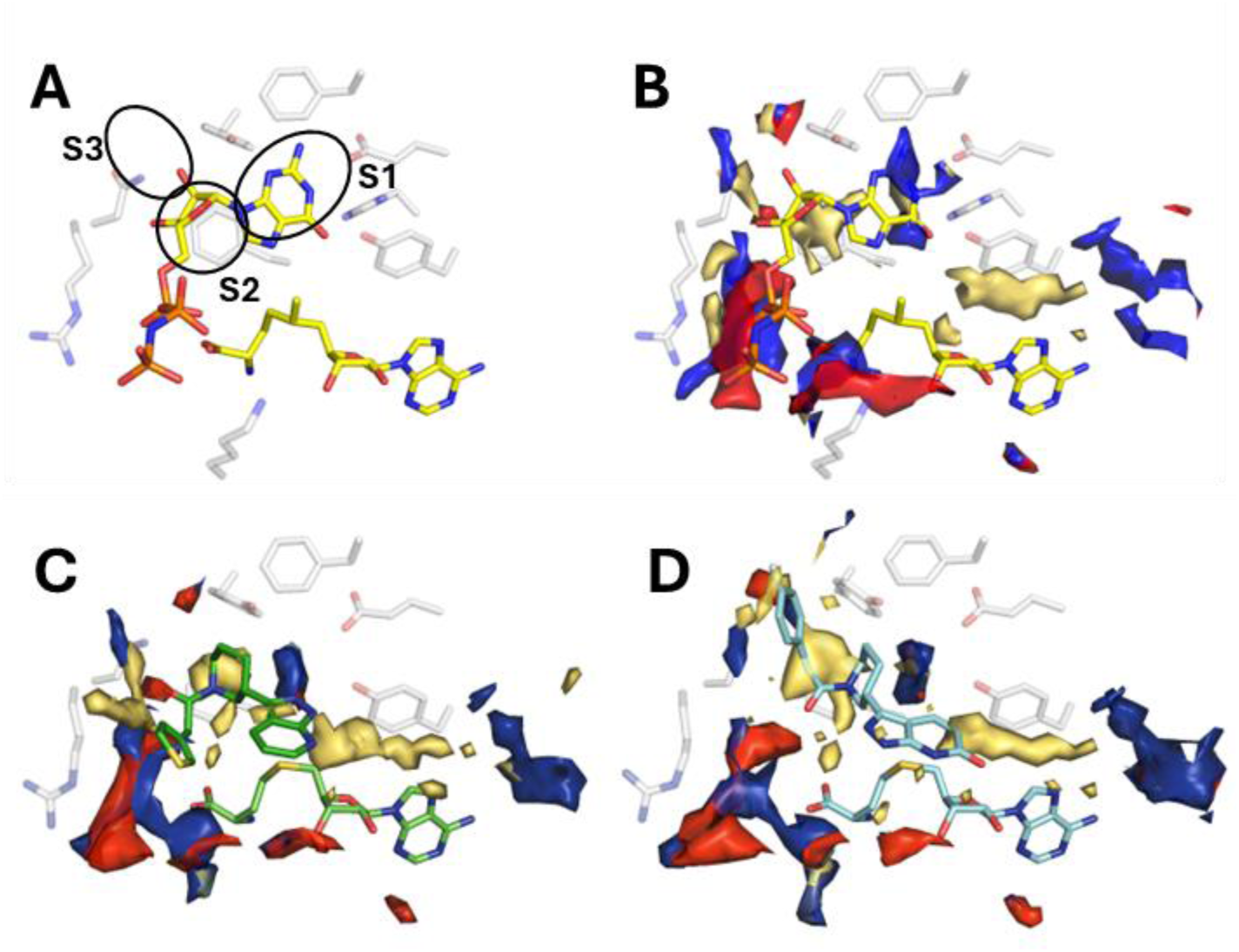
Potential compound development routes informed by MIF-based structures. The GRID MIFs calculated for the RNMT polypeptide chain (with corresponding energy cut-offs in brackets) are contoured yellow (hydrophobic: −1.5 kcal/mol), blue (hydrogen-bond donor: −5.5 kcal/mol) and red (hydrogen-bond acceptor: −5.0 kcal/mol). (A) sinefungin, (B) cap analogue with amine modified to methyl group, (C) SAH-DDD1060606 and (D) SAH-DDD1870799 crystal structures.

The biophysical characterisation of cap binding to Avi-*Hs*RNMT-RAM described earlier, in combination with the observed conformation of the guanine group, indicate that the steric effect of the amine on the accompanying sinefungin/methyl (SAM) group in our crystal structures are important factors for stabilisation. It is only when the sinefungin or SAM is in place that the interactions observed in the cap structure (polar with H288, Y289, E370, Y467 and Pi stacking with F285) become important. Thus, in the presence of SAH, any new inhibitor ligand should ideally carry features that take into account the steric effect of the methyl on SAM to permit the guanine-mimicking group access to the S1 pocket.

## Discussion

Despite a growing interest in *Hs*RNMT-RAM as a valuable target to either inhibit the proliferation and increase the apoptosis of cancer cells [14,16,20,21], or to prevent eukaryotic virus proliferation [33], the lack of understanding of the enzymatic mechanism limits the development of selective and efficient drugs. Here we present a comprehensive overview of the enzymatic mechanism of HsRNMT monomer and in complex with its co-factor RAM. This characterisation is used as a basis to discuss *Hs*RNMT-RAM druggability through the description of two selective inhibitors targeting the cap pocket.

In our manuscript, we describe a label-free RapidFire mass spectrometry assay for the enzymatic characterization of *Hs*RNMT-RAM. Although our data are robust, we observed significant differences compared to the recently published data [34]. Specifically, there were 10- and 40-fold differences in the K_m_ values for SAM and RNA, respectively. There are many potential sources for these differences. These include detail variation in the expression systems for the protein, the presence or absence of RAM and divergence in the presence or absence of a small RNA attached to the 3’ end of the cap substrate. Despite these differences we report comparable IC_50_ values for sinefungin, which differ by less than 2-fold.

Biophysical characterisation of substrate binding shows an ordered reaction with SAM required to bind first to enable subsequent cap binding. While an ordered reaction for RNMT has been made [10], the observed order of substrate addition is reversed to the prediction.

We report the structure of two small molecule inhibitors of RNMT as result of a diversity driven hit discovery campaign. These molecules behave biochemically as uncompetitive inhibitors with respect to both cap and SAM despite the structural biology data leading to an expectation of competitive inhibition with cap. The biophysical data resolves this puzzle by demonstrating that the compounds require the presence of the enzymatic product SAH in RNMT to bind efficiently. This stabilisation of enzyme-product complex explains the competitive binding resulting in uncompetitive inhibition. The model we propose for the behaviour of RNMT in binding its substrates and the compounds is presented (Figure 9). This is similar to the mode of action of mycophenolic acid against the inosine 5’ monophosphate dehydrogenase [35,36].

**Figure 9.**
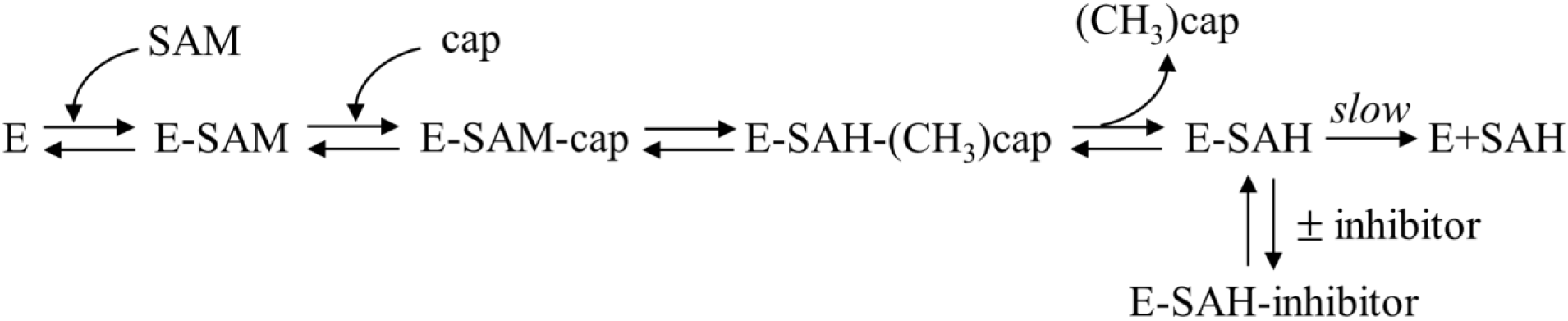
Diagram summarising the ordered bi-bi mechanism of enzymatic activity of RNMT and the enzyme-product stabilisation driven by binding of the compounds resulting in the observed uncompetitive inhibition.

Finally, we used molecular modelling approaches to map the physicochemical properties of the cap pocket of HsRNMT. This, in combination with the crystal co-complexes of the enzyme with the small molecule hits, allows us prediction of chemical modifications that will generate better tool molecules and potentially new medicines.

## Materials & Methods

Reagents were purchased from Sigma Aldrich (Gillingham, Dorset, UK) unless otherwise stated.

### Protein Production

For biochemical assays, full length *Hs*RNMT (1-476) was cloned in the pOPC bi-cistronic plasmid as a TEV-cleavable His-MBP fusion with *Hs*RAM (1-90) and expressed in *E. coli* BL21 (DE3) using LB medium. This protein is referred to as *Hs*RNMT-RAM.

For SPR, full length *Hs*RNMT was cloned into a proprietary pNIC28a with an N-terminal Avi-tag and *Hs*RAM (1-90) into a pACYC vector encoding a C-terminal TEV cleavable hexa-His tag and is described as Avi-*Hs*RNMT-RAM in this manuscript. Plasmids were co-transformed into *E.coli* BL21 (DE3) in presence of a pCDFduet-BirA vector for in-vivo biotinylation and grown in TB autoinduction media. For crystallography, the *Hs*RNMT catalytic domain (165-476, Δ416-455 GLGC) was cloned into pET15bTEV and expressed as a TEV-cleavable His tag fusion in *E. coli* BL21 DE3 using ZY autoinduction medium.

Both *Hs*RNMT full length proteins (1-476) in complex with *Hs*RAM (1-90) (*Hs*RNMT-RAM and Avi-*Hs*RNMT-RAM) were purified using the same procedure. Harvested cells were re-suspended in a lysis buffer comprising 20mM HEPES, 175mM NaCl, 25mM KCl, 10% glycerol, 0.5mM TCEP, pH 7.5 supplemented with DNAse and EDTA-free protease inhibitors (Sigma) before lysis at 30 psi with a cell disruptor (Constant Systems). *Hs*RNMT-RAM was initially purified by Ni-NTA affinity chromatography (HisTrap FF 5 ml, Cytiva) in the same buffer and eluted with an imidazole gradient. Imidazole was then quickly removed from the fractions of interest using a HiPrep 26/10 desalting column in the same buffer. The desalted fractions were later incubated with the TEV protease (1:20) overnight at 4°C prior to a second purification by Ni affinity chromatography performed in the same conditions. The flowthrough, containing the target protein, was concentrated and purified further with gel filtration chromatography on a Superdex 75 26/60, pre-equilibrated with the same buffer. Monomeric fractions were pooled and concentrated to 15mg/ml before snapshot freezing in liquid nitrogen. The purification of *Hs*RNMT catalytic domain (165-476) Δ416-455 GLGC was purified in a similar way with the following buffer 20mM HEPES, 350mM NaCl, 50mM KCl, 10 % glycerol, 0.5mM TCEP, pH 7.5.

### MTase-Glo Screen

A selection of 48,806 diverse drug like compounds from our internal compound library were tested for inhibition against *Hs*RMNT-RAM at a single concentration, using MTase-Glo, a bioluminescent based assay from Promega (Madison, WI, USA). Briefly, compounds were incubated with 5 nM *Hs*RMNT-RAM, 2µM SAM and 2µM cap, using buffer conditions specified in the MTase-Glo assay kit, for 60 minutes. Following this incubation, the MTase-Glo Reagent and Detection solution (at ratios of 5:2 and 1:1 of the final assay volume, respectively) were added to each well and the signal allowed to develop for a further 30 minutes in the dark. The resulting luminescence was read on a PHERAstar FS (BMG). A hit was defined as percent effect was greater than mean + 3 standard deviations of the screen results (50.8%). Following this screen, compounds that had been identified as hits were confirmed by retesting at 10-point dose response in the RapidFire assay. Potency was calculated by fitting data to a 4-parameter logistical fit (model 203) using Sigmaplot Version 14.0.3.192. Data are presented as the mean of 3 independent experiments with 95% Confidence intervals (95%CI).

### RapidFire Assay

An endpoint 384-well plate assay for *Hs*RMNT-RAM activity was developed using the Agilent RapidFire 365 high-throughput system with integrated SPE interfaced with the Agilent 6740 triple quadrupole mass spectrometer. The assay was performed by mixing 5nM *Hs*RNMT-RAM in buffer (20mM Tris, pH 8.0, 50mM NaCl) supplemented with 1mM EDTA, 1mM DTT, 0.1mg/mL BSA, 0.005% Nonidet P40 (Roche) and 3mM MgCl2 with 2µM of substrates (G(5’)ppp(5’)G Sodium Salt (cap) [New England Biolabs, Ipswich, MA]), and S-(5′ Adenosyl)-L-methionine chloride dihydrochloride (SAM [Cayman Chemical, Ann Arbor, MI]). The enzymatic reaction incubated for 60 min at room temperature followed by quenching with 80 μL 1% formic acid (VWR, Radnor, PA) containing 0.03 µg/µL S-adenosylhomocysteine-d4 (d4SAH [Cambridge Bioscience, Cambridge, UK]). Plates were centrifuged at 500×g for 1 min after every addition and sealed using the PlateLoc thermal microplate sealer before analysis on the RapidFire. Samples were loaded onto the RapidFire system and analysis performed as described for the identification of Inhibitors of the SARS-CoV-2 Guanine-N7-Methyltransferase [37].

### K_m_ Determinations

Experiments to determine the K_m_^app^ values were carried out using a 2-fold dilution of cap or SAM from 40 - 0.08µM and a saturating concentration (40µM) of SAM (K_m_^app^ cap) or cap (K_m_^app^ SAM). *Hs*RMNT-RAM was used at 5nM. Data were analysed to calculate the initial velocity (V0). K_m_^app^ and V_max_ values were determined by nonlinear regression analysis fitted to the Michaelis–Menten equation [38] using Sigmaplot Version 14.0.3.192. Data are presented as the mean of 3 independent experiments with 95%CI.

### Mode of Inhibition Experiments

Mode of inhibition experiments were carried out by testing enzymatic activity at a range of compound and substrate concentrations. Compounds were tested at 5 different concentrations selected around the pIC_50_ against 5 substrate concentrations (between 2.2 and 20µM). The data were fitted globally to equations describing inhibition [39] to determine the respective kinetic constants. All analyses were performed in Sigmaplot Version 14.0.3.192.

Best fit was determined using AICc (Akaike information criterion corrected [40], as well as reference to the standard error of the fit [41]. Where relevant, data are presented as the mean of 3 independent experiments with 95%CI.

### Surface plasmon resonance (SPR)

Studies of binding kinetics were performed on a Biacore™ 8K+ instrument (GE Healthcare). Full length *Hs*RNMT complexed to *Hs*RAM 1-90 (Avi-*Hs*RNMT-RAM) was immobilized in HBS-P+ buffer (10 mM HEPES, 150 mM NaCl, 0.5mM TCEP, 0.05% BSA and 0.05% Tween 20) on a Series S high-affinity streptavidin (SA) sensor chip by C-terminal Avi-tag capture, to an immobilization level of 3300 RU. Compounds were tested in duplicate a in three-fold, ten-point serial dilution series in immobilization buffer supplemented with 1% DMSO, with a flow rate of 30 µl min^-1^, a contact time of 60s and a dissociation time of 120s.

All monitored binding resonance signals were referenced with responses from unmodified reference surfaces, corrected for bulk shifts arising from differences in DMSO concentrations between samples and running buffer (solvent correction) and blank-referenced. Analysis is performed using the Biacore™ Insight Evaluation Software 2.0 with data fitted using the Langmuir 1:1 model.

### Isothermal titration calorimetry (ITC)

Full length *Hs*RNMT-RAM was dialysed overnight at 4°C against 25 mM HEPES pH 7.5, 150 mM NaCl, 1 mM TCEP supplemented with 10μM sinefungin or 10μM SAH. Working stock for ligands and protein complex was prepared at the required concentration by diluting the stocks with the dialysis buffer. The DMSO concentrations in the cell and syringe solutions were matched to 2.5% for the compound (DDD1060606 or DDD1870799) titrations. Experiments were performed using 13 injections of 3 μl of ligand using Malvern PEAQ-ITC system at 25°C, at 750rpm with reference power set to 7μcal/s. The thermograms were analysed using MicroCal PEAQ-ITC Analysis Software, blanked to ligand control, and fit to One Set of Sites Model.

### Crystallisation and structure determination

*Hs*RNMT (165-476) Δ416-455 GLGC (*Hs*RNMT) was buffer exchanged in PIPES 20mM, NaCl 200mM, glycerol 10%v/v, TCEP 1mM, pH 6.5 using Zeba 10 kDa desalting columns (Thermofisher) and concentrated to 17mg/ml. Co-complexes structures were achieved by mixing SAH (1mM) with the appropriate inhibitor (5mM) whereas the crystallization of cap was only possible by a combination of sinefungin (1mM) with GMP-PnP 10mM and 2mM MgCl_2._ *Hs*RNMT was crystallized at 17°C using the sitting drop vapor diffusion method by mixing 200 nl of the protein solution mixed with 200 nl of a reservoir solution containing 0.1M MES pH 6.3-6.9, 50 mM Na_2_SO_4_, 18-25% PEG 6000. Crystals grew in 48 hours and were flash-frozen in liquid nitrogen in the reservoir solution supplemented with 28% glycerol. Data sets were collected at 100K at beamline synchrotron ESRF-ID23 or using a Rigaku M007HF copper-anode generator fitted with Varimax Cu-VHF optics and a Saturn 944HG+ CCD detector. Data were processed using XDS [42] in P21 and scaled using Aimless [43] to a resolution between 2.0-2.5Å. The following structure solution and refinement activities were performed. The initial phases were determined by molecular replacement using Phaser [44] from the CCP4 suite [45] using the human catalytic domain *Hs*RNMT (pdb code 5E8J) with *Hs*RAM removed and the segment 416-455 as the search model. The initial density maps were further improved by solvent flattening and histogram matching using RESOLVE [47] as implemented in the Phenix suite [48]. The model was refined through iterative cycles of model building and computational refinement with COOT [49] and Phenix Refine (Phenix suite). Data measurements and model refinement statistics are presented in Table S1. Coordinate files and associated experimental data have been deposited in the Protein Data Bank (PDB) with accession codes 8Q9W, 8Q69 and 8Q8G.

## Author contributions

**Lesley-Anne Pearson**: Conceptualisation, Validation, Methodology, Formal analysis, Investigation, Data Curation, Writing – Original Draft, Visualisation**. Alain-Pierre Petit**: Conceptualisation, Validation, Methodology, Formal analysis, Investigation, Data Curation, Writing – Original Draft, Visualisation**. Cesar Mendoza Martinez**: Investigation, Writing – Original Draft, **Fiona Bellany**; Resources, **De Lin**: Methodology, Investigation, **Sarah Niven**: Investigation, **Rachel Swift**: Investigation, **Thomas Eadsforth**: Resources, **Paul Fyfe**: Writing – Original Draft, Resources, **Marilyn Paul**; Investigation, **Vincent Postis**; Conceptualisation, Supervision, **Xiao Hu**: Formal Analysis, **Victoria H. Cowling:** Conceptualisation, Writing – Original Draft, Writing – Review and Editing, **David W Gray**: Conceptualisation, Validation, Data Curation, Writing – Original Draft, Writing – Review and Editing, Supervision, Funding acquisition.

## Acknowledgements

The authors recognises the valuable input from our Compound Management (Alex Cookson, Steve Bell, Kirsty Cookson, Fraser Hughes) and Data Management (Steve Thompson, Edan Gardner, Aren Lai, Kashish Sharma) Teams.

## Data availability statement

All crystallographic data are deposited in the Protein Data Bank. All other data are contained within the manuscript.

