## supplementary data for "Characterisation of RNA guanine-7 methyltransferase (RNMT) using a small molecule approach"

### Slide 1
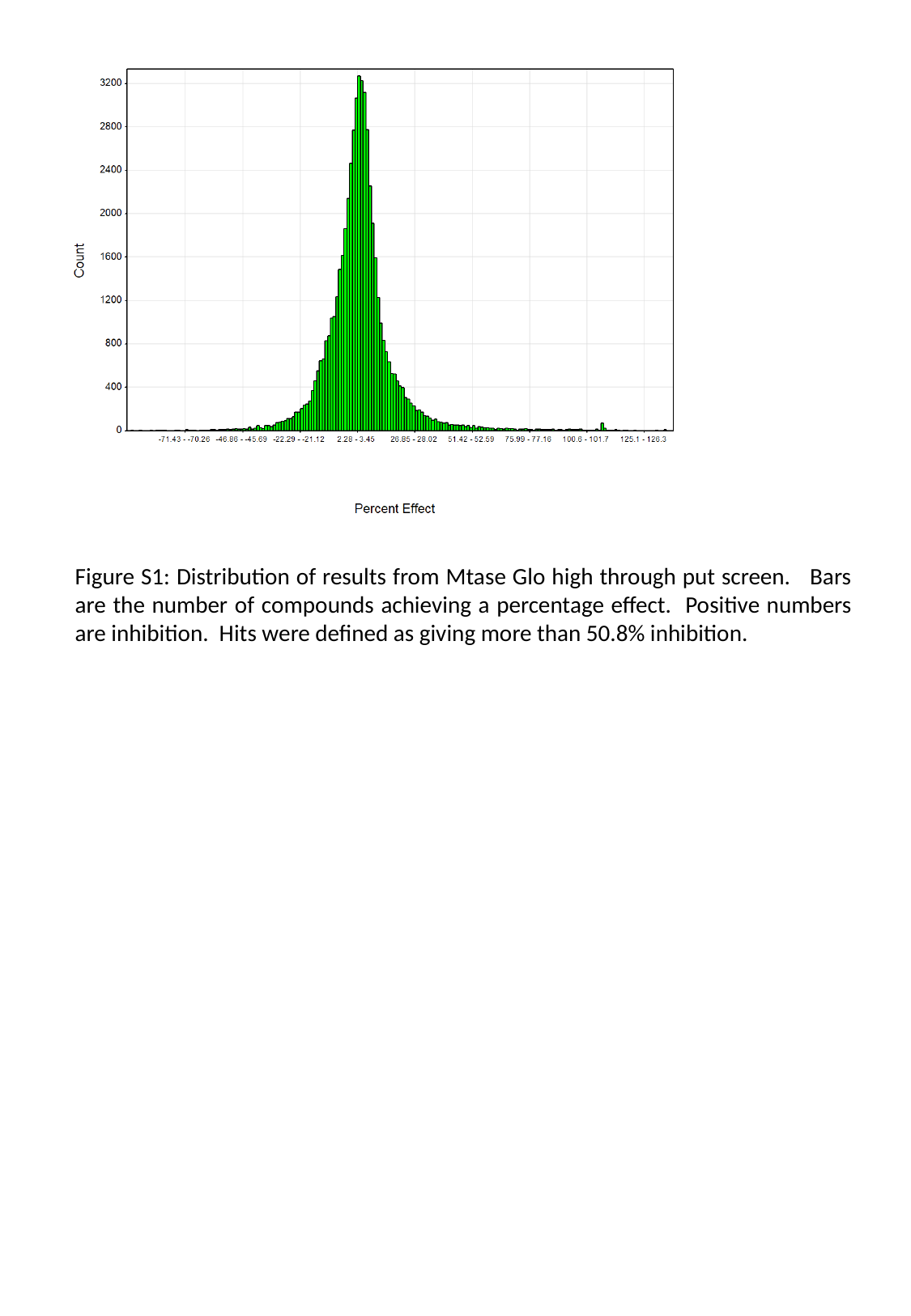

Figure S1: Distribution of results from Mtase Glo high through put screen. Bars are the number of compounds achieving a percentage effect. Positive numbers are inhibition. Hits were defined as giving more than 50.8% inhibition.

### Slide 2
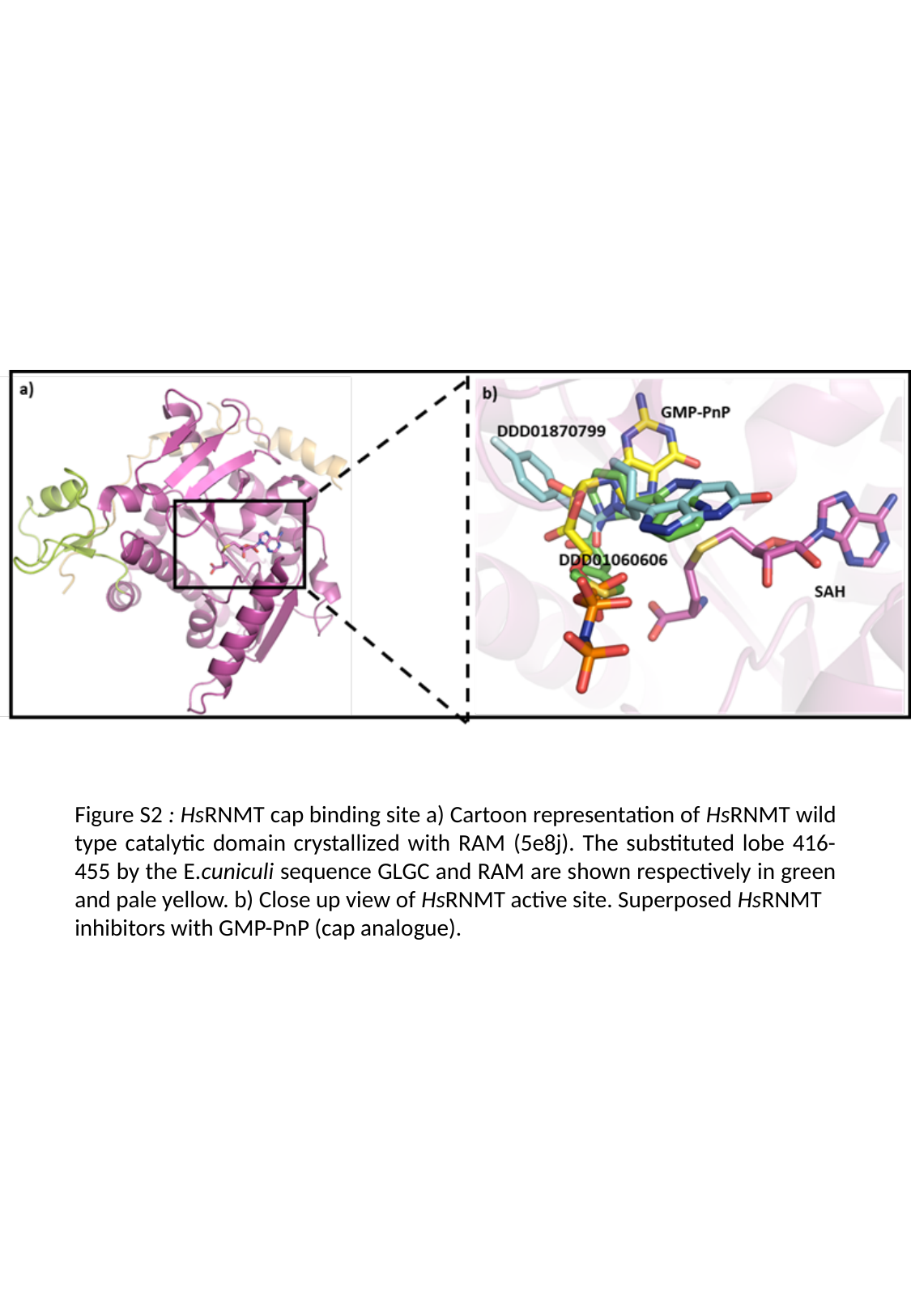

Figure S2 : HsRNMT cap binding site a) Cartoon representation of HsRNMT wild type catalytic domain crystallized with RAM (5e8j). The substituted lobe 416-455 by the E.cuniculi sequence GLGC and RAM are shown respectively in green and pale yellow. b) Close up view of HsRNMT active site. Superposed HsRNMT
inhibitors with GMP-PnP (cap analogue).

### Slide 3
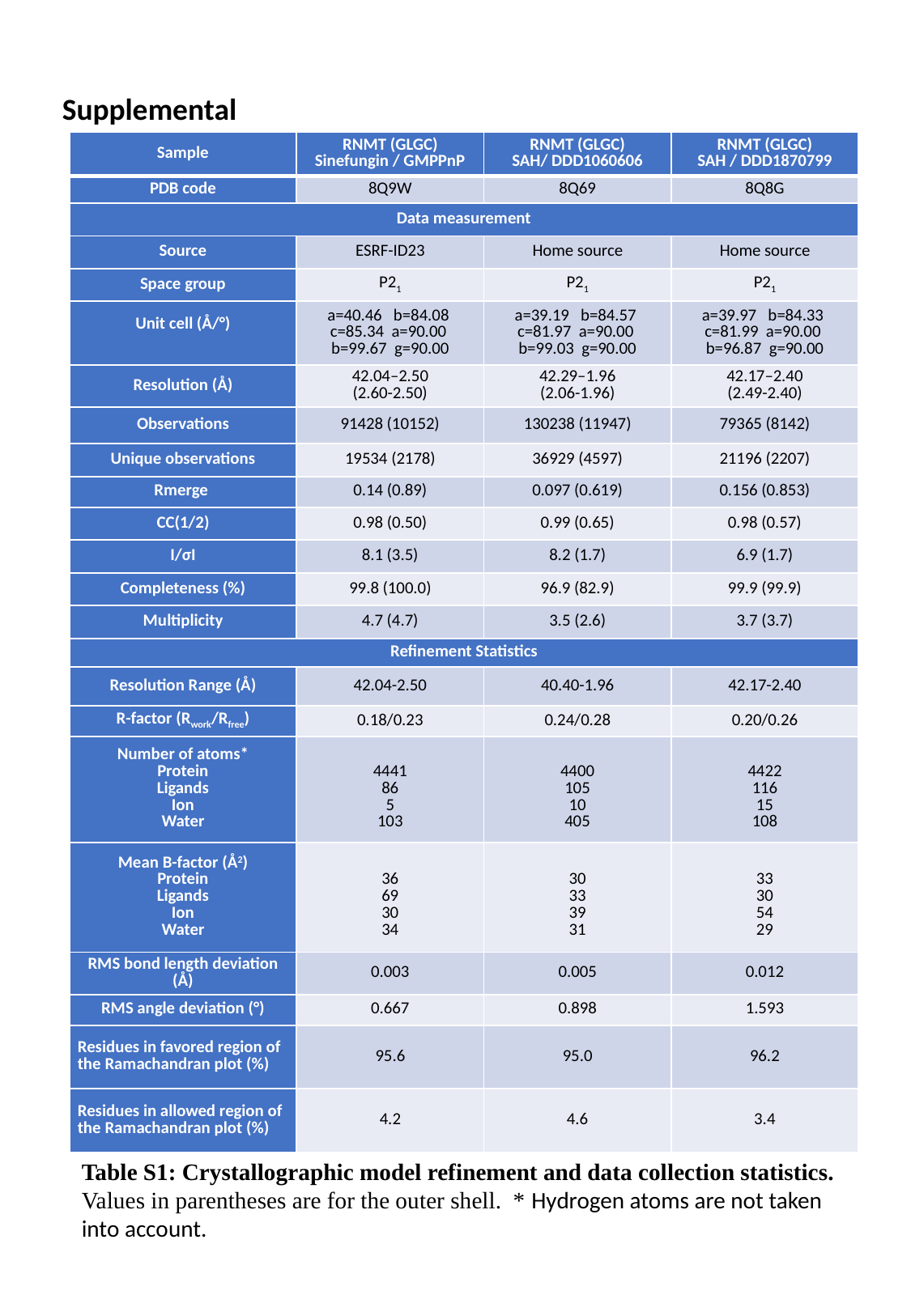

Supplemental
| Sample | RNMT (GLGC) Sinefungin / GMPPnP | RNMT (GLGC) SAH/ DDD1060606 | RNMT (GLGC) SAH / DDD1870799 |
| --- | --- | --- | --- |
| PDB code | 8Q9W | 8Q69 | 8Q8G |
| Data measurement | | | |
| Source | ESRF-ID23 | Home source | Home source |
| Space group | P2_1_ | P2_1_ | P2_1_ |
| Unit cell (Å/°) | a=40.46 b=84.08 c=85.34 a=90.00 b=99.67 g=90.00 | a=39.19 b=84.57 c=81.97 a=90.00 b=99.03 g=90.00 | a=39.97 b=84.33 c=81.99 a=90.00 b=96.87 g=90.00 |
| Resolution (Å) | 42.04–2.50 (2.60-2.50) | 42.29–1.96 (2.06-1.96) | 42.17–2.40 (2.49-2.40) |
| Observations | 91428 (10152) | 130238 (11947) | 79365 (8142) |
| Unique observations | 19534 (2178) | 36929 (4597) | 21196 (2207) |
| Rmerge | 0.14 (0.89) | 0.097 (0.619) | 0.156 (0.853) |
| CC(1/2) | 0.98 (0.50) | 0.99 (0.65) | 0.98 (0.57) |
| I/σI | 8.1 (3.5) | 8.2 (1.7) | 6.9 (1.7) |
| Completeness (%) | 99.8 (100.0) | 96.9 (82.9) | 99.9 (99.9) |
| Multiplicity | 4.7 (4.7) | 3.5 (2.6) | 3.7 (3.7) |
| Refinement Statistics | | | |
| Resolution Range (Å) | 42.04-2.50 | 40.40-1.96 | 42.17-2.40 |
| R-factor (R_work_/R_free_) | 0.18/0.23 | 0.24/0.28 | 0.20/0.26 |
| Number of atoms\* Protein Ligands Ion Water | 4441 86 5 103 | 4400 105 10 405 | 4422 116 15 108 |
| Mean B-factor (Å^2^) Protein Ligands Ion Water | 36 69 30 34 | 30 33 39 31 | 33 30 54 29 |
| RMS bond length deviation (Å) | 0.003 | 0.005 | 0.012 |
| RMS angle deviation (°) | 0.667 | 0.898 | 1.593 |
| Residues in favored region of the Ramachandran plot (%) | 95.6 | 95.0 | 96.2 |
| Residues in allowed region of the Ramachandran plot (%) | 4.2 | 4.6 | 3.4 |
Table S1: Crystallographic model refinement and data collection statistics. Values in parentheses are for the outer shell. * Hydrogen atoms are not taken into account.

### Slide 4
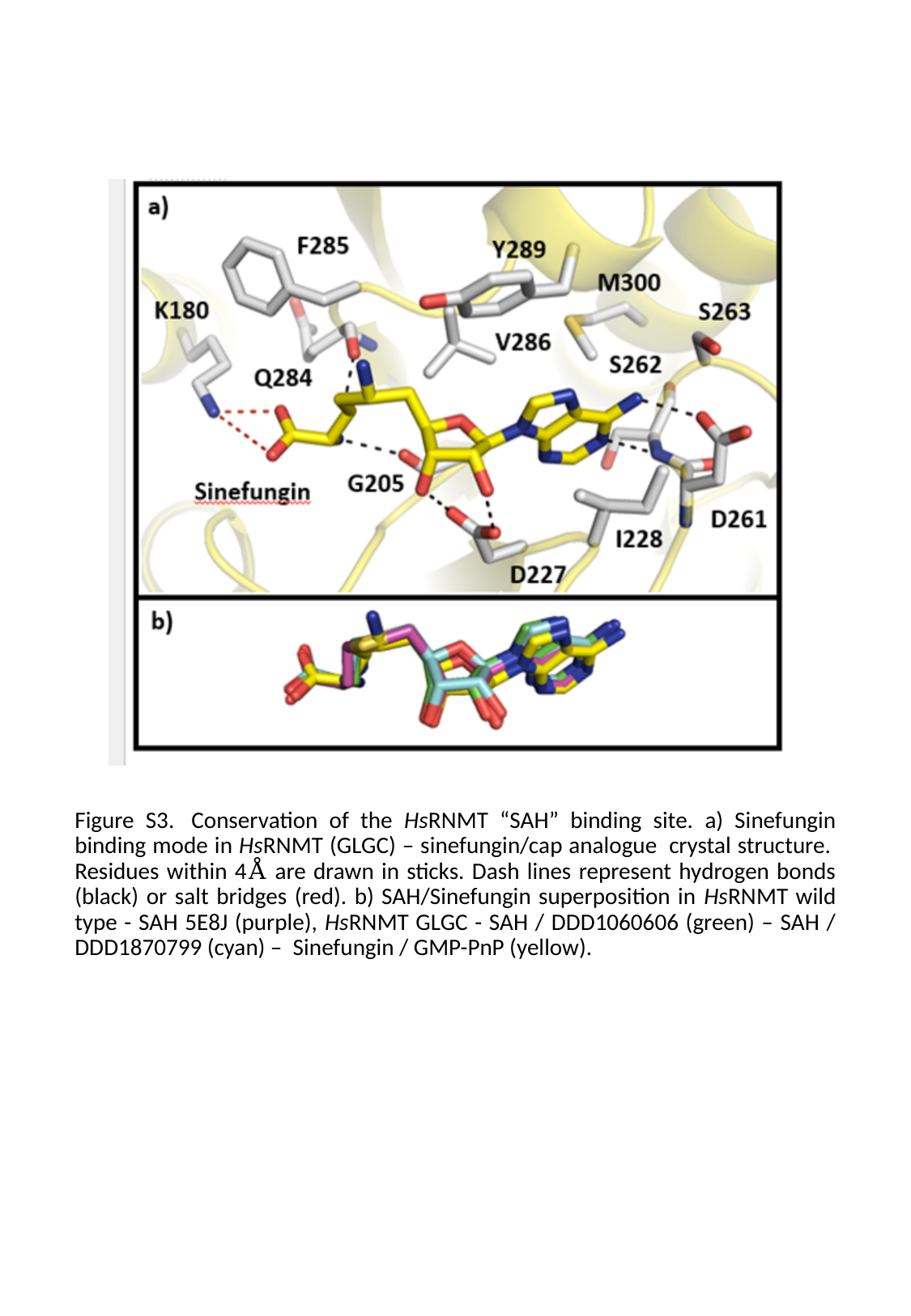

Figure S3.  Conservation of the HsRNMT “SAH” binding site. a) Sinefungin binding mode in HsRNMT (GLGC) – sinefungin/cap analogue  crystal structure.  Residues within 4Å are drawn in sticks. Dash lines represent hydrogen bonds (black) or salt bridges (red). b) SAH/Sinefungin superposition in HsRNMT wild type - SAH 5E8J (purple), HsRNMT GLGC - SAH / DDD1060606 (green) – SAH / DDD1870799 (cyan) –  Sinefungin / GMP-PnP (yellow).

### Slide 5
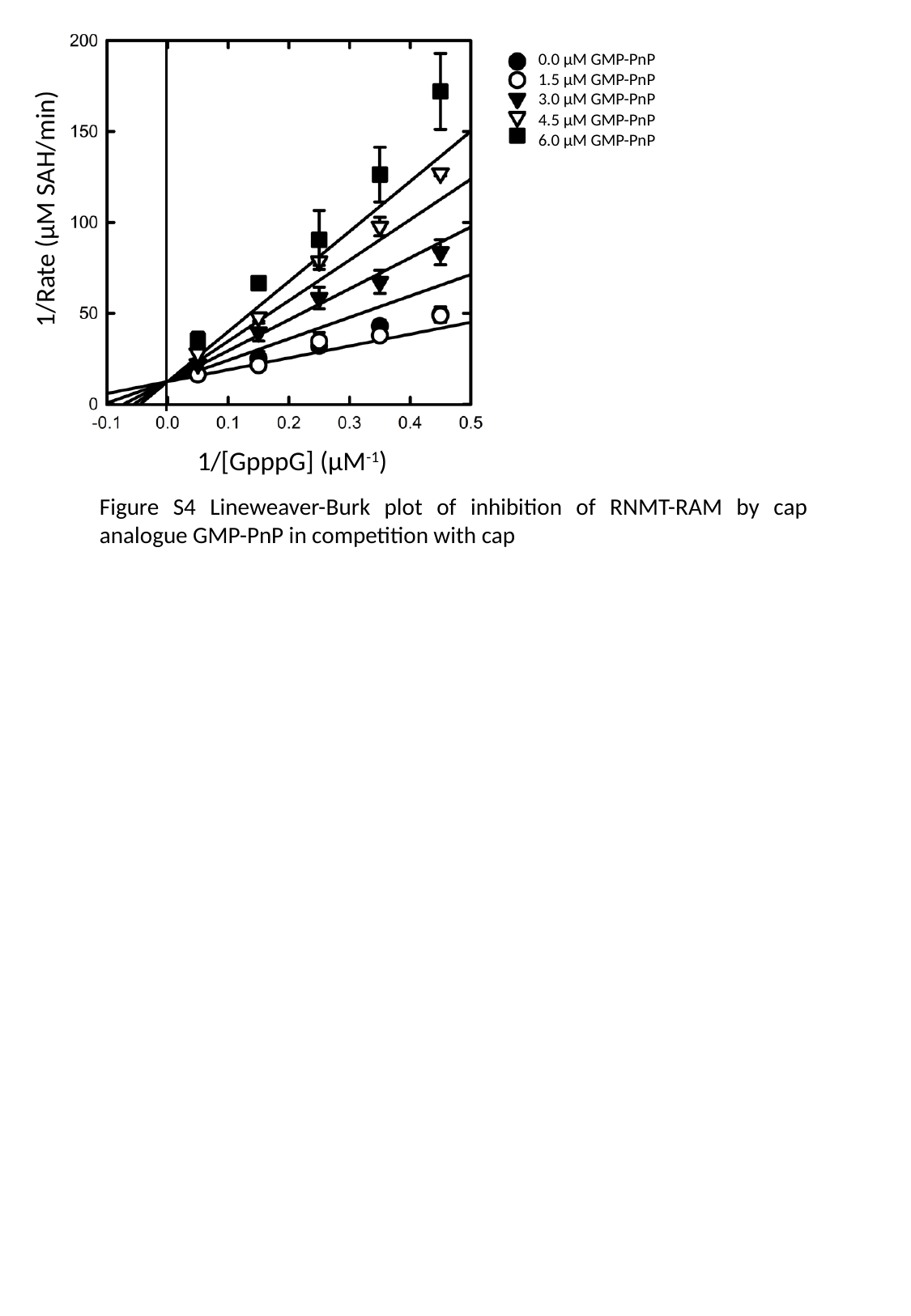

0.0 µM GMP-PnP
1.5 µM GMP-PnP
3.0 µM GMP-PnP
4.5 µM GMP-PnP
6.0 µM GMP-PnP
uM
uM
uM
uM
uM
1/Rate (µM SAH/min)
1/[GpppG] (µM^-1^)
Figure S4 Lineweaver-Burk plot of inhibition of RNMT-RAM by cap analogue GMP-PnP in competition with cap

### Slide 6
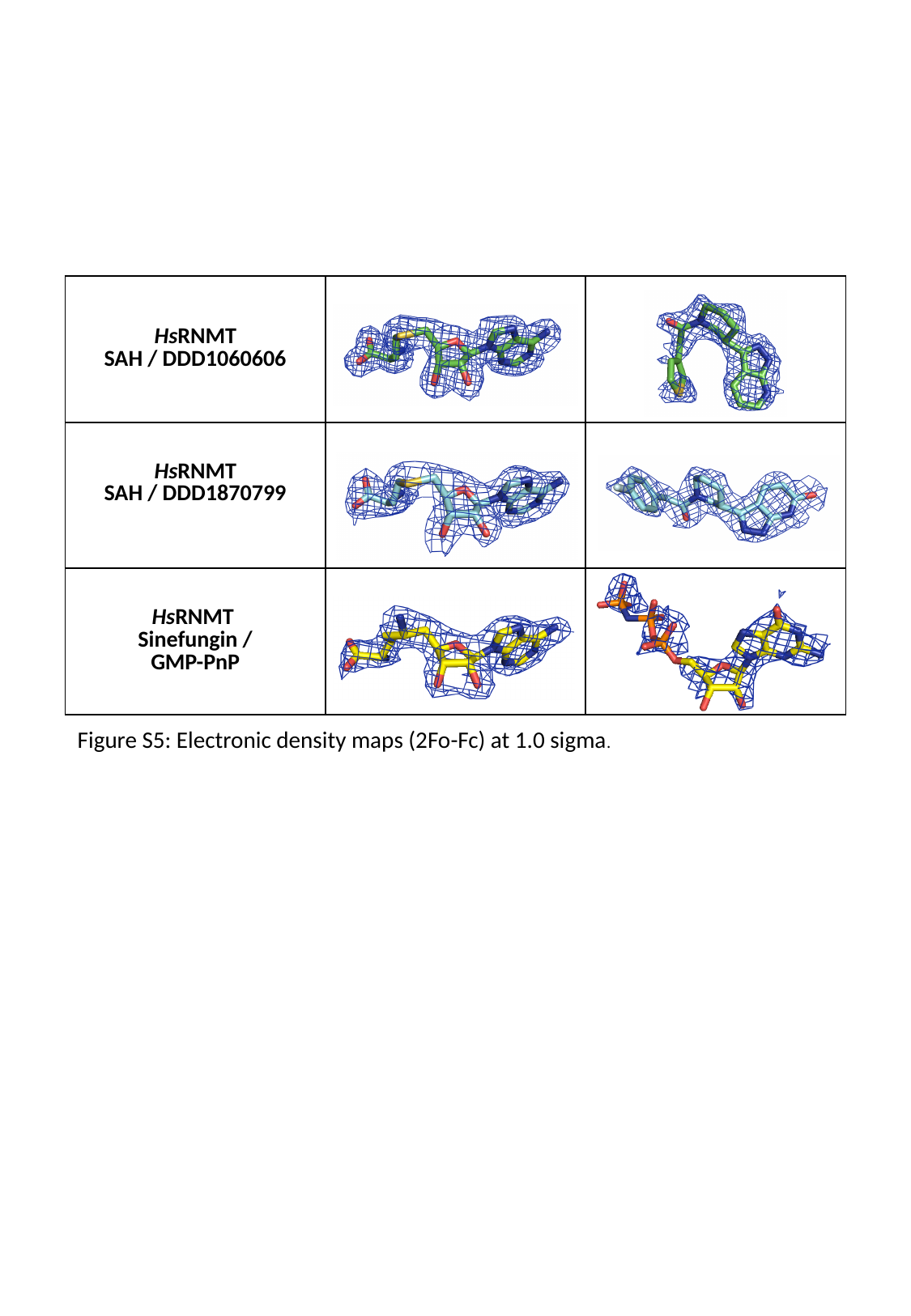

| HsRNMT SAH / DDD1060606 |
| --- |
| HsRNMT SAH / DDD1870799 |
| HsRNMT Sinefungin / GMP-PnP |
Figure S5: Electronic density maps (2Fo-Fc) at 1.0 sigma.
